# Modulation of Intrauterine Adhesion Fibrosis by Lactobacillus delbrueckii: Insights from ERK1/2 Pathway Inhibition and Multi-omics Analyses

**DOI:** 10.64898/2026.02.15.706062

**Authors:** Bao Liu, Jing Xu, Mingqian Chen, Dongzhi Gou, Yugang Chi

## Abstract

**Background:** Intrauterine adhesion (IUA) is an important cause of infertility and menstrual disorders, substantially reducing quality of life and increasing healthcare costs. Current treatments, including hysteroscopic adhesiolysis and hormonal therapy, show limited efficacy and high recurrence, highlighting the need for new therapeutic strategies.

**Methods:** We evaluated the antifibrotic effects of *Lactobacillus delbrueckii* in IUA, with a focus on ERK1/2 signaling inhibition. A multimodal framework integrating clinical outcomes, cellular and animal experiments, uterine microbiome profiling (16S rRNA sequencing), transcriptomics, and metabolomics was used to characterize host–microbe–metabolite interactions relevant to fibrosis.

**Results:** Clinically, patients receiving *L. delbrueckii* showed lower adhesion formation scores(AFS) and reduced recurrence. Microbiome profiling demonstrated increased uterine microbial diversity and altered community composition in treated patients, accompanied by reduced expression of fibrosis-associated markers. In both cellular and mouse models, *L. delbrueckii* decreased fibrotic marker expression and reduced ERK1/2 phosphorylation. Metabolomics identified elevated 3-hydroxyanthranilic acid (3-HAA) in treated groups, suggesting a potential role in immune and inflammatory modulation.

**Conclusion:** *L. delbrueckii* attenuates IUA-associated fibrosis, at least in part, by suppressing ERK1/2 phosphorylation and increasing the abundance of the metabolite 3-HAA. These findings support the therapeutic potential of *L. delbrueckii* for improving uterine repair and reproductive outcomes. Future studies should validate efficacy and safety in larger, diverse cohorts and assess long-term outcomes.

## 1. Introduction

Intrauterine adhesions (IUA), also referred to as Asherman’s syndrome, are defined by fibrous bands within the uterine cavity that arise after endometrial trauma, infection, or intrauterine procedures^[1]^. IUA represent a persistent clinical challenge because they can cause hypomenorrhea or amenorrhea, infertility, recurrent pregnancy loss, and pregnancy complications, with substantial consequences for quality of life and healthcare utilization^[2]^. The core pathology involves damage to the basal endometrium and disruption of normal regenerative programs. In this setting, wound-healing responses become maladaptive, characterized by fibroblast activation, excessive extracellular matrix deposition, and scar-like remodeling that distorts uterine cavity architecture^[3]^. The rising incidence of IUA underscores the need for improved recognition, diagnosis, and treatment.

The current standard of care—hysteroscopic adhesiolysis followed by hormonal therapy to support endometrial regrowth—improves uterine cavity patency in many patients, yet recurrence remains common, especially in moderate-to-severe disease^[4]^. The variability of outcomes underscores two key limitations: (i) insufficient suppression of the fibrotic cascade that drives re-adhesion, and (ii) incomplete restoration of functional endometrium despite anatomical separation. Consequently, there is strong interest in strategies that target fibrosis and inflammation while improving the uterine microenvironment for regeneration^[5]^.

A growing literature implicates the uterine and vaginal microbiota in reproductive health, implantation, and gynecologic disorders. Lactobacillus-dominant communities in the lower reproductive tract are generally associated with favorable outcomes, whereas dysbiosis can promote inflammation and compromise epithelial and stromal homeostasis. Probiotic approaches that restore beneficial taxa may therefore provide a noninvasive method to modify local immune tone and repair responses^[6]^. However, for IUA, most evidence remains indirect: while probiotics can reduce inflammatory mediators and influence mucosal immunity, the mechanistic links between probiotic exposure and fibrotic remodeling of endometrium have not been systematically established.

At the signaling level, the mitogen-activated protein kinase (MAPK) network—and particularly the ERK1/2 cascade—controls cell proliferation, differentiation, stress responses, and matrix remodeling^[7]^. Aberrant ERK1/2 activation has been reported in multiple fibrotic disorders, where it can promote myofibroblast differentiation, enhance production of TGF-β–responsive genes, and increase collagen synthesis^[8]^. While the MAPK family also includes p38 and JNK pathways, the relative dominance of these branches varies by tissue and injury context. In the uterus, how ERK1/2 activity integrates inflammatory cues and wound-healing programs during adhesion formation is not well defined. Moreover, an emerging concept is that microbial metabolites—particularly tryptophan catabolites—can regulate immune and stromal cell signaling, providing a biochemical bridge between microbiome composition and host tissue remodeling^[9]^.

In this study, we tested the hypothesis that *Lactobacillus delbrueckii* attenuates endometrial fibrosis associated with IUA by suppressing ERK1/2 activation and by increasing production or accumulation of antifibrotic metabolites. We combined (i) a randomized clinical intervention with vaginal *L. delbrueckii* after hysteroscopic adhesiolysis, (ii) multi-omics profiling of uterine secretions (16S rRNA microbiome sequencing and LC–MS/MS metabolomics), and (iii) mechanistic validation in cellular and mouse models using transcriptomics and pharmacologic perturbation of ERK1/2 signaling. This integrated design enables causal inference across levels—microbiome composition, metabolite changes, signaling activity, and fibrosis-associated phenotypes—and provides a translational framework for probiotic-based approaches to prevent IUA recurrence.

## 2. Materials and methods

### 2.1 Study design, participant enrollment, and specimen collection

All clinical procedures were conducted in accordance with the Declaration of Helsinki and were approved by the Ethics Committee of Chongqing Health Center for Women and Children (Approval No. 2021044). Ninety patients diagnosed with IUA and treated by hysteroscopic adhesiolysis at Chongqing Maternal and Child Health Hospital between January and August 2023 were enrolled.

During surgery, endometrial secretions were collected using a cold-knife technique under sterile conditions. After adhesiolysis, a uterine balloon was placed for 5 days as part of postoperative management. Participants were randomized into two groups: a control group (n = 45) receiving sequential estrogen–progestogen therapy, and an intervention group receiving the same hormonal regimen plus commercial *L. delbrueckii* administered vaginally for 10 days (Lac group; n = 45). During treatment, four participants missed doses and one withdrew; thus, 40 participants were included in the final Lac group analysis. One month after surgery, endometrial secretions were collected again for 16S rRNA sequencing and metabolomics. Patients underwent hysteroscopic re-evaluation, and pregnancy status was recorded at 1 year postoperatively.

Inclusion criteria were age 18–45 years, no recent medication use, and no major preoperative abnormalities. Exclusion criteria included estrogen-related diseases and severe systemic illness.

### 2.2 Uterine microbiome profiling by 16S rRNA sequencing

Microbial genomic DNA was extracted from uterine lavage samples using the FastPure Stool DNA Isolation Kit. DNA quality and concentration were assessed by agarose gel electrophoresis and NanoDrop spectrophotometry. Samples were stored at −80°C until analysis.

The V3–V4 hypervariable regions of the bacterial 16S rRNA gene were amplified using primers 338F and 806R on a T100 Thermal Cycler under standardized PCR conditions. Amplicons were quantified, purified, and sequenced on an Illumina NextSeq 2000 platform. After quality filtering, amplicon sequence variants (ASVs) were inferred, and taxonomic annotation was performed using QIIME2. Rarefaction curves and rank-abundance curves were generated to assess sequencing depth and coverage. Alpha-diversity indices (e.g., Chao, Shannon, Simpson) and beta-diversity analyses (NMDS and PCoA) were computed to compare microbial community richness and structure between groups.

### 2.3 Metabolomics profiling of uterine secretions

Samples were stored at −80°C prior to metabolomics. Approximately 50 mg of each sample was transferred to 2mL tubes. A methanol–water extraction solution containing an internal standard was added, and samples were homogenized with a 6 mm bead using a cryogenic grinder (−10°C, 50 Hz, 6 min). Samples underwent ultrasonic extraction (5°C, 30 min) and were centrifuged at 13,000 g for 15 min at 4°C. Supernatants were transferred to vials for LC–MS/MS analysis. All processing was performed using consistent volumes and under sterile conditions to minimize contamination.

Quality control (QC) samples were used to assess instrument stability. Relative standard deviation (RSD) and cumulative peak proportion were evaluated to confirm data suitability for downstream analyses. Differential metabolites were identified using multivariate models (PLS-DA and OPLS-DA) and univariate statistics, and pathway enrichment was conducted using KEGG. Random forest modeling with recursive feature elimination and cross-validation (RFECV) was applied to prioritize candidate biomarkers; receiver operating characteristic (ROC) curves were used to quantify discriminative performance.

### 2.4 Cell culture and establishment of an LPS-induced fibrotic model

Immortalized human endometrial stromal cells (IHESCs; CTCC-001-0425), STR-verified and purchased from Zhejiang Meisen Cell Technology Co., Ltd., were cultured in DMEM/F12 (Thermo Fisher Scientific) supplemented with 10% fetal bovine serum and 1% penicillin/streptomycin (Invitrogen) at 37°C in 5% CO₂.

To establish a fibrotic phenotype, IHESCs were seeded in 6-well plates and cultured for 48 h. After adherence, cells were washed with PBS and treated with lipopolysaccharide (LPS; 2 μg/mL) for 48 h. Morphological changes were recorded. Fibrosis-associated markers (TGF-β1, α-SMA, collagen I) were quantified by RT-qPCR, immunofluorescence, and/or western blotting.

### 2.5 Culture of *Lactobacillus delbrueckii* and treatment of fibrotic IHESCs

Live *Lactobacillus delbrueckii* (DM8909) powder was obtained from Inner Mongolia Shuangqi Pharmaceutical Co., Ltd. Powder (0.2–0.5 g) was cultured in 200 mL LBS medium anaerobically at 37°C for 18 h. Cultures were streaked onto LBS agar and incubated anaerobically at 37°C for 48 h. Colonies were transferred to LBS broth and cultured until the fermentation broth reached pH 4.7–4.8. Cultures were further incubated with shaking for 24–48 h. Bacterial counts were determined to prepare defined inocula.

For cell experiments, groups included NC, LPS, and LPS + *L. delbrueckii* at varying concentrations (CFU/mL). After LPS induction, IHESCs were co-cultured with *L. delbrueckii* for 3 h. Cells were then processed for measurement of fibrosis markers (TGF-β1, α-SMA, collagen I) and MAPK pathway proteins as appropriate.

To test whether secreted metabolites mediated effects, cell cultures were treated with defined proportions of *L. delbrueckii* culture supernatant (after removal of bacteria) and compared with direct bacterial addition.

### 2.5 Mouse IUA model and probiotic treatment

All animal procedures were approved by the institutional ethics committee (Approval No. IACUC-CQMU-2024-0627). Female SPF-grade C57 mice (8–10 weeks) were obtained from the Experimental Animal Center of Chongqing Medical University.

IUA was induced by mechanical endometrial injury combined with LPS exposure. Briefly, under anesthesia and sterile conditions, a 1 mL syringe needle was bent and used to scrape the uterine wall repeatedly. Bleeding at the incision site was used as confirmation of effective injury. Sutures pre-soaked in LPS solution were then placed into the uterine cavity to promote adhesion formation. Control mice underwent anesthesia and laparotomy with suturing to maintain procedural consistency.

For probiotic exposure, *L. delbrueckii* was administered vaginally once daily at 1 × 10¹⁰ CFU/mL (1 mL) for 7 days. For mechanistic experiments, mice were assigned to five groups (n = 10/group): control, IUA, IUA + LAC, IUA + ERK1/2 inhibitor, and IUA + ERK1/2 agonist + LAC. The ERK1/2 inhibitor was administered intraperitoneally at 1 mg/kg daily; the ERK1/2 agonist was administered intraperitoneally at 10 mg/kg daily. Vaginal LAC dosing remained 1 × 10¹⁰ CFU/mL (1 mL) daily.

### 2.6 Immunofluorescence Staining Protocol for Cells

Coverslips were cleaned and soaked in 75% ethanol for 10 minutes, then heated to remove ethanol before being placed in a six-well plate to cool. Endometrial stromal cells (IHESCs) were digested with trypsin, resuspended, counted, and seeded at 1-2 × 10⁴ cells/mL. After 6–12 hours, cells were washed with PBS and treated with fresh medium or 2 μg/mL LPS for 48 hours. Post-incubation, cells were fixed with 4% paraformaldehyde for 10–20 minutes and permeabilized with a permeabilization buffer. Blocking was done with 3% hydrogen peroxide, followed by 3% BSA or serum. Primary antibody incubation occurred overnight at 4°C, followed by washing and incubation with HRP-conjugated secondary antibody. Detection was achieved using TSA and TBST washing. Antibody elution was performed with incubation at 37°C for 30 minutes, followed by further washing. For nuclear staining, DAPI was applied for 10 minutes, and cells were mounted with antifade medium. Images were captured using appropriate filters for fluorophore detection.

### 2.7 Western blotting

Total protein was extracted, and concentrations were determined using a BCA Protein Assay Kit (Nanjing Jiancheng Bioengineering Institute). Membranes were blocked with 5% skim milk for 2 h at room temperature and washed three times with TBST (10 min each). Membranes were incubated with primary antibodies overnight at 4°C, washed with TBST, and incubated with HRP-conjugated anti-rabbit secondary antibodies for 1 h at room temperature. After three additional TBST washes, bands were visualized using a Bio-Rad Gel Doc EZ Imager, and densitometry was performed with ImageJ.

### 2.8 Immunohistochemistry

Excised tissues were rinsed with saline, fixed in formalin, embedded in paraffin, and sectioned. Sections were dewaxed in xylene and rehydrated through graded ethanol. Endogenous peroxidase was blocked with 3% H₂O₂ for 30 min. Antigen retrieval was performed in boiling sodium citrate buffer (0.01 M, pH 6.0) for 10 min. After blocking with goat serum for 30 min, sections were incubated overnight in a humidified chamber with primary antibodies against TGF-β1 (21898-1-AP, 1:100; Proteintech) and p-ERK (ET1610-13, 1:100; Hua An). Sections were incubated with secondary antibodies (Proteintech) for 1 h at room temperature and developed with DAB. Negative controls omitted primary antibodies (PBS substituted). Two pathologists independently evaluated staining intensity and classified expression as high, low, or negative.

### 2.9 Statistical Analysis

SPSS 22.0 and R (v4.2.2) were used for statistical analyses. Normally distributed continuous variables were compared using a two-sample t-test (two groups) or one-way ANOVA (multiple groups). GraphPad Prism and ImageJ were used for visualization and densitometry. A two-sided P < 0.05 was considered statistically significant.

## 3. Results

### 3.1 Clinical efficacy

Baseline characteristics were comparable between groups. Mean age was 32.05 ± 5.84 years in the Lactobacillus group and 30.93 ± 5.28 years in the control group (P = 0.36). The number of curettage procedures did not differ (2.78 ± 1.90 vs. 2.69 ± 2.03; P = 0.84). Baseline adhesion formation score (AFS) was also similar (7.18 ± 1.97 vs. 6.62 ± 2.35; P = 0.25). Adhesion etiology, menstrual status, and estrogen dose showed no significant between-group differences (P = 0.22, 0.19, and 0.34, respectively; **Supplementary Table 1**). Biochemical indicators were likewise comparable (**Supplementary Table** 2). Although postoperative menstrual recovery and pregnancy rates did not differ significantly, the Lactobacillus group exhibited greater improvement in AFS, suggesting a potential reproductive benefit that warrants further mechanistic and long-term investigation (**Supplementary Table 3**).

### 3.2 Microbial diversity and community structure before and after surgery

Preoperatively, uterine microbial diversity did not differ between groups. Alpha-diversity indices (Chao and Shannon) and beta-diversity analyses showed no significant differences (P > 0.05; **Fig. 1A–D**), and community composition at phylum and genus levels was broadly similar (**Fig. 1E–F**). These findings support baseline comparability prior to intervention.

**Figure 1.**
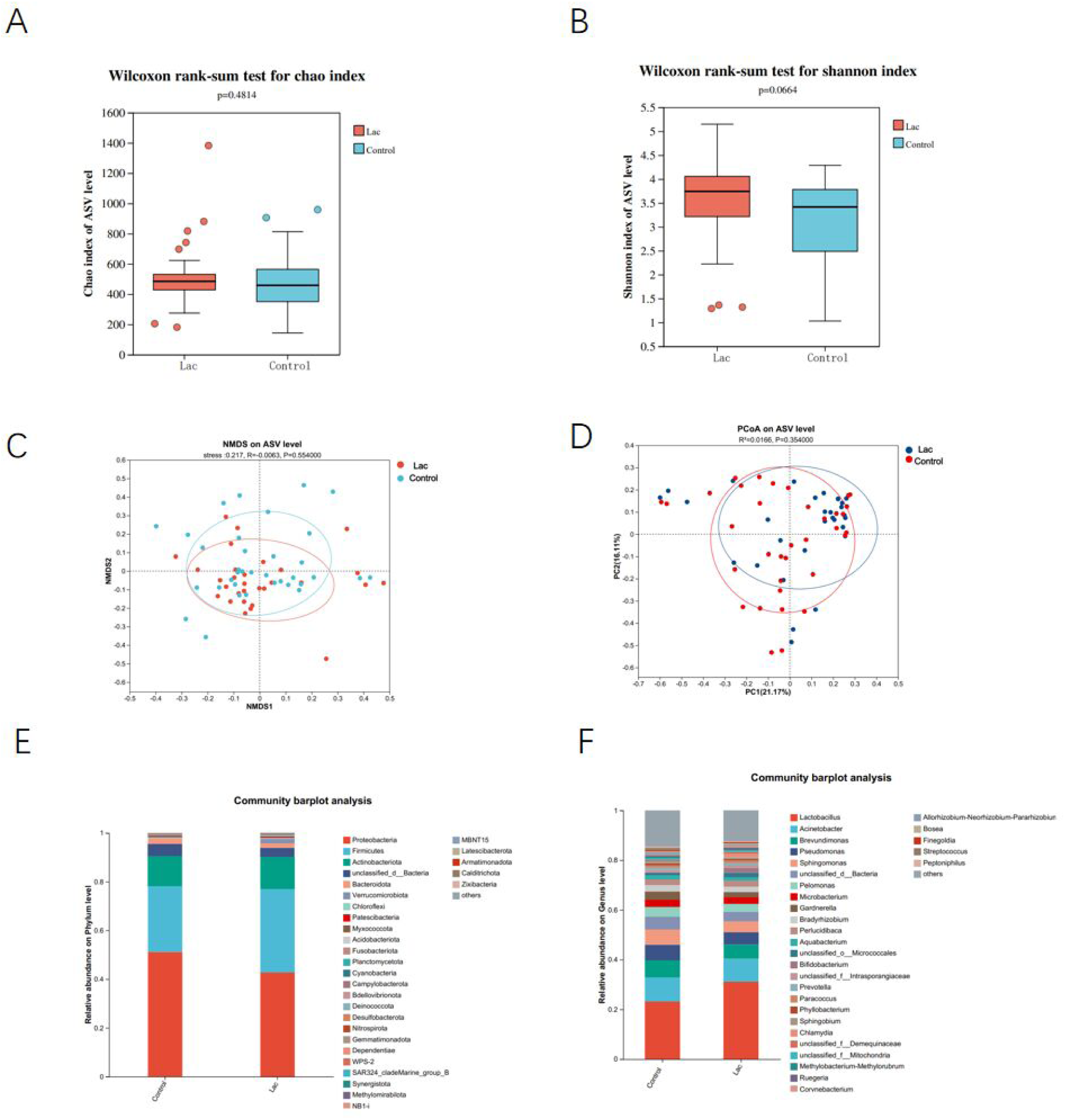
Preoperative microbial diversity and community structure. (A) Chao index. (B) Shannon index. (C) NMDS. (D) PCoA. (E) Phylum-level relative abundance. (F) Genus-level relative abundance.

Postoperatively, alpha-diversity differed significantly between groups. Simpson and Shannon indices indicated higher diversity and richness in the Lac group than in controls (P < 0.05; **Fig. 2A–B**). NMDS and PCoA analyses demonstrated significant separation between groups (P < 0.05; **Fig. 2C–D**), indicating that vaginal administration of *L. delbrueckii* substantially altered community structure in the uterine cavity.

**Figure 2.**
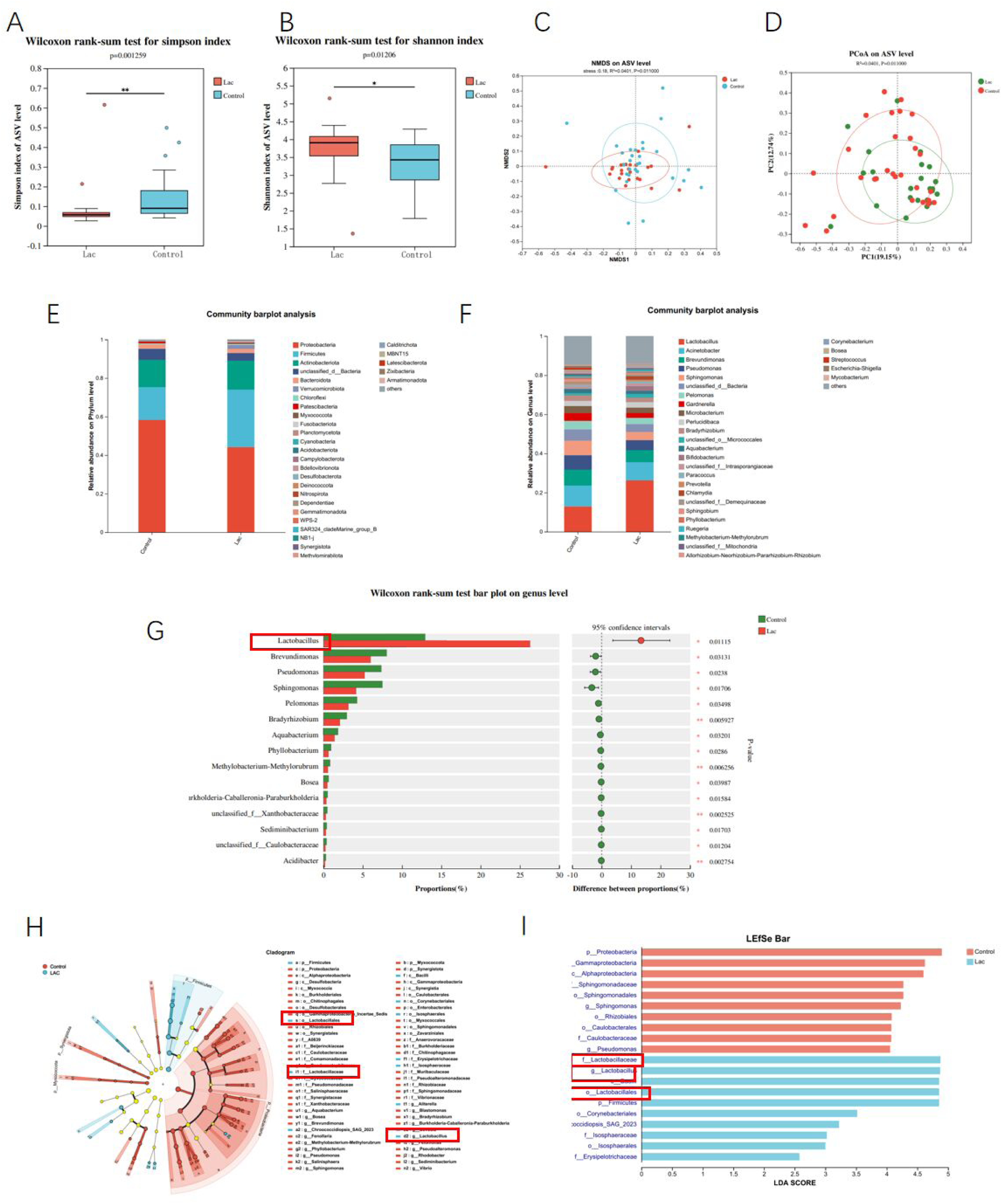
Postoperative microbial diversity and community structure. (A) Simpson index. (B) Shannon index. (C) NMDS. (D) PCoA. (E) Phylum-level relative abundance. (F) Genus-level relative abundance. (G) Differential genera between groups. (H) LEfSe cladogram. (I) LDA scores.

At the phylum level, both groups remained dominated by Proteobacteria, Firmicutes, and Actinobacteriota. At the genus level, the most abundant taxa included *Lactobacillus*, *Acinetobacter*, *Brevundimonas*, and *Pseudomonas*. Despite similarities in dominant taxa, relative abundances differed between groups; *Lactobacillus*, *Brevundimonas*, and *Pseudomonas* showed significant between-group differences (**Fig. 2E–G**).

To identify taxa driving these differences, LEfSe analysis was performed across multiple taxonomic ranks. The results revealed distinct discriminative features in each group and highlighted strong enrichment of *Lactobacillus* in the Lac group (**Fig. 2H–I**), consistent with successful colonization or enrichment following probiotic administration.

### 3.3 Metabolomics analysis

Quality control indicated good analytical stability: the cumulative proportion of peaks exceeded 0.9, meeting criteria for downstream analysis (**Fig. 3A**). From the mass spectrometry dataset, 4273 metabolites were detected, of which 309 were differentially abundant between groups (**Fig. 3B**). Compared with controls, 102 metabolites were upregulated and 207 were downregulated in the Lactobacillus group. KEGG pathway enrichment suggested that differential metabolites were primarily involved in metabolic pathways (**Fig. 3C–D**).

**Figure 3.**
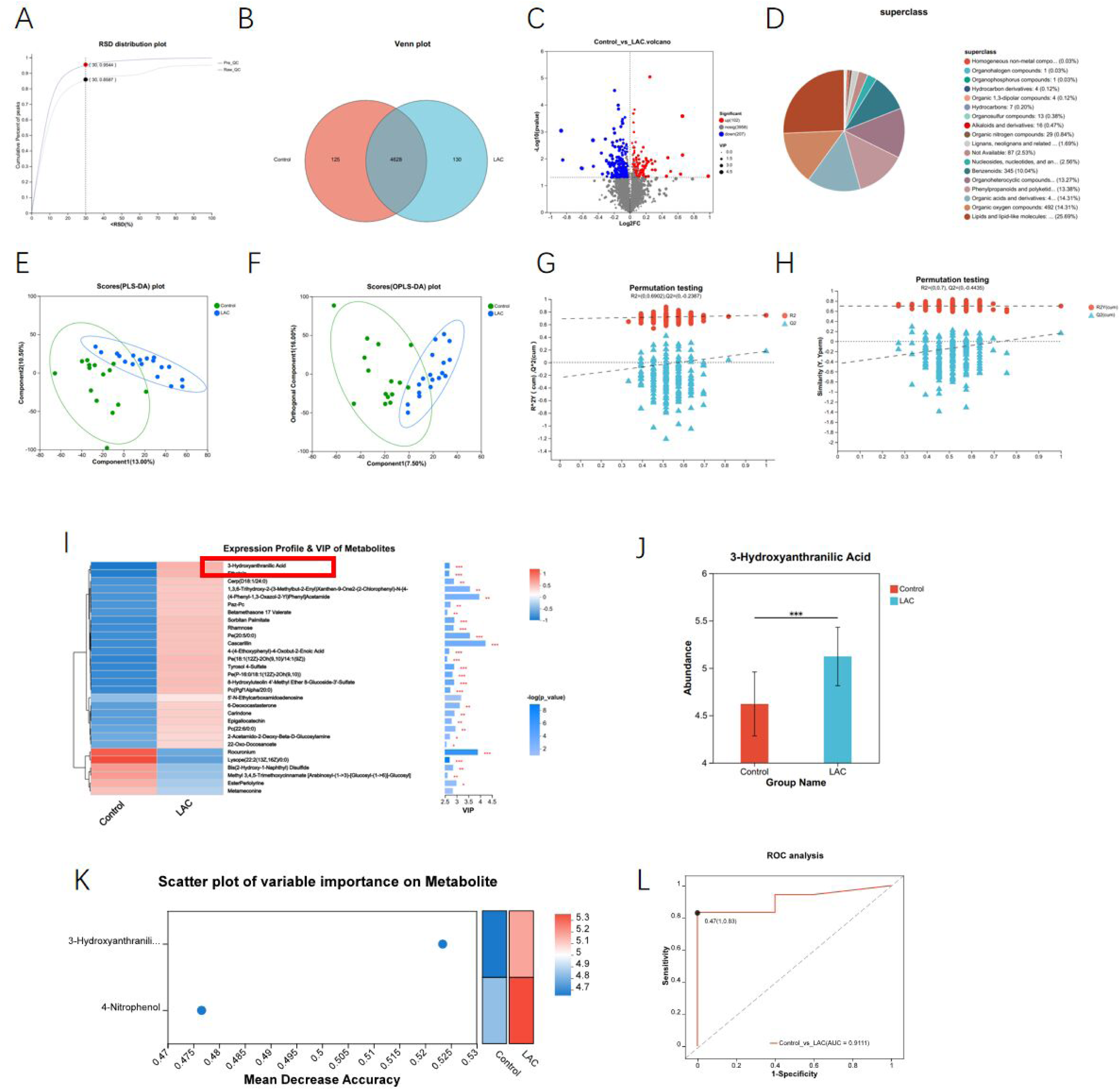
Metabolomics profiling of uterine secretions. (A) RSD plot. (B) Venn diagram. (C) Volcano plot. (D) Compound classification. (E) PLS-DA score plot. (F) OPLS-DA score plot. (G) PLS-DA permutation test (200 iterations). (H) OPLS-DA permutation test (200 iterations). (I) VIP ranking. (J) Differential abundance of 3-HAA. (K) Feature-importance ranking. (L) ROC curve.

PLS-DA and OPLS-DA analyses identified clear separation between groups. Metabolites with VIP > 1 and P < 0.05 were defined as significant discriminators (**Fig. 3E–F**). Model robustness was supported by 200 permutation tests, in which the Q2 intercepts were below zero for both models, indicating no evidence of overfitting (**Fig. 3G–H**). These findings suggest substantial biological differences in metabolic states between groups.

VIP-based ranking from the OPLS-DA model highlighted 3-hydroxyanthranilic acid (3-HAA) as markedly increased in the Lactobacillus group. Consistently, 3-HAA showed pronounced between-group differences, implicating it as a candidate mediator associated with endometrial fibrosis regulation (**Fig. 3I–J**).

To prioritize biomarkers, random forest modeling with recursive feature elimination and cross-validation (RFECV) was applied. Feature importance ranking identified 3-HAA as the top candidate biomarker. ROC analysis yielded an AUC of 0.91, indicating strong classification performance (**Fig. 3K–L**).

### 3.4 MAPK signaling in the fibrotic cell model

Following establishment of the LPS-induced fibrosis model, *Lactobacillus* was added at varying concentrations for 3h. Western blotting showed the most pronounced reduction of fibrosis-associated proteins at 1 × 10^7^ CFU/mL (**Fig. 4A**), indicating a dose-dependent antifibrotic effect of *Lactobacillus delbrueckii*. To investigate underlying mechanisms, transcriptomic profiling was performed in IHESCs from the LPS and LPS + *Lactobacillus* groups, followed by differential gene analysis and KEGG enrichment (**Fig. 4B–C**). Differentially expressed genes were significantly enriched in focal adhesion, cytokine–cytokine receptor interaction, MAPK signaling, steroid biosynthesis, and mannose-type O-glycan biosynthesis pathways (all P < 0.05; **Fig. 4D–E**), implicating MAPK signaling as a candidate regulatory axis.

**Figure 4.**
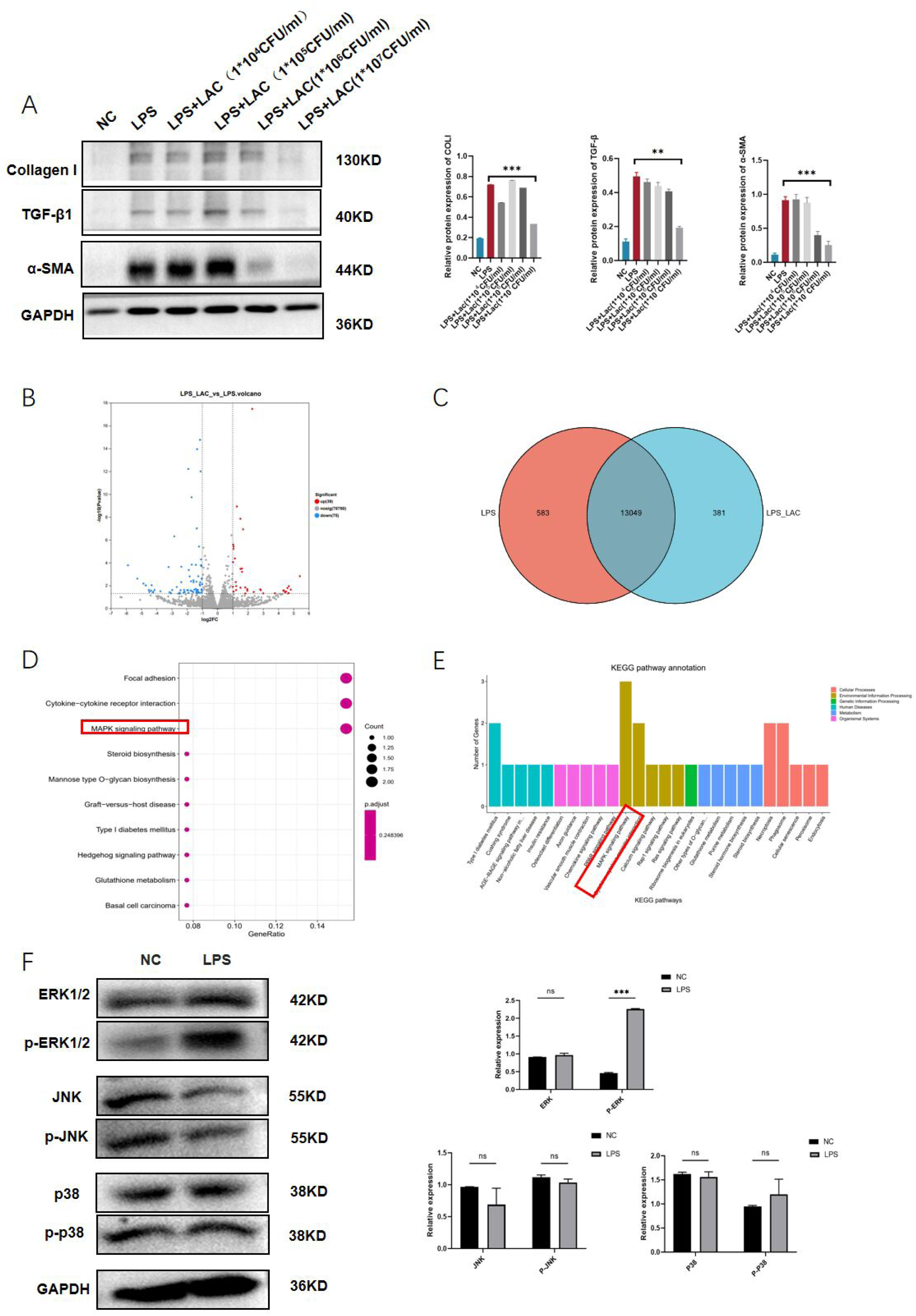
Pathway screening and MAPK signaling in the cellular fibrosis model. (A) Effects of L. delbrueckii concentration on fibrosis markers in IHESCs. (B) Volcano plot of transcriptomics. (C) Venn diagram. (D) KEGG enrichment dot plot. (E) KEGG enrichment bar plot. (F) MAPK pathway protein expression in the fibrosis model.

MAPK signaling comprises three major subfamilies: ERK1/2, p38, and JNK. Western blotting showed that p-ERK1/2 was significantly increased in the LPS group relative to controls (P < 0.05), whereas total ERK1/2 and the p38/JNK pathways (p38, p-p38, JNK, p-JNK) did not differ significantly (P > 0.05; **Fig. 4F**). These results suggest selective activation of ERK1/2 phosphorylation in this model and support ERK1/2 as a potential target of *L. delbrueckii*.

### 3.5 *Lactobacillus delbrueckii* suppresses fibrosis through ERK1/2 signaling

To validate ERK1/2 involvement in the antifibrotic effect of *Lactobacillus*, an ERK1/2 agonist was introduced. Five groups were assessed: NC, LPS, ERK1/2 agonist, LPS + LAC, and LPS + ERK1/2 agonist + LAC. Immunofluorescence and western blotting showed that ERK1/2 agonist treatment significantly increased p-ERK1/2, accompanied by elevated fibrosis markers (TGF-β1, α-SMA, and collagen I). In contrast, addition of *Lactobacillus* reduced p-ERK1/2 and downregulated TGF-β1 and collagen I compared with both the agonist and LPS groups (P < 0.05; **Fig. 5A–B**).

**Figure 5.**
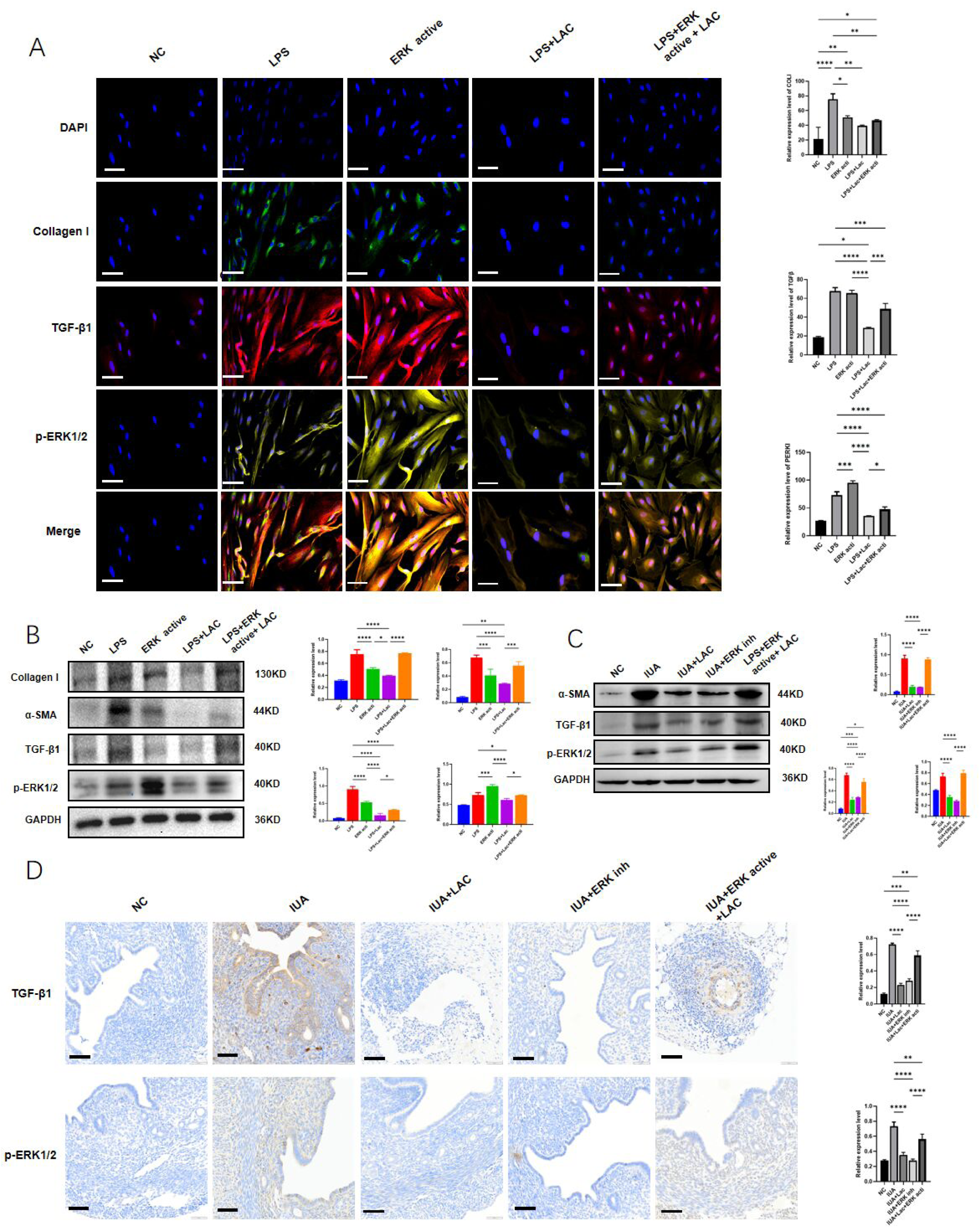
Functional validation of ERK1/2 involvement in vitro and in vivo. (A) Immunofluorescence of TGF-β1, collagen I, and p-ERK1/2 after ERK1/2 agonist exposure (scale bar = 20 μm). (B) Western blot of TGF-β1, collagen I, α-SMA, and p-ERK1/2. (C) Western blot of uterine TGF-β1, α-SMA, and p-ERK1/2 in mouse groups. (D) Immunohistochemistry of TGF-β1 and p-ERK1/2 (scale bar = 50 μm).

In vivo, mice were randomized into five groups (n = 10/group): control, IUA model, IUA + LAC, IUA + ERK1/2 inhibitor, and IUA + ERK1/2 agonist + LAC. After model establishment, the ERK1/2 inhibitor was administered intraperitoneally daily (1 mg/kg). In the agonist + LAC group, the ERK1/2 agonist was administered intraperitoneally daily (10 mg/kg), and LAC was administered vaginally daily (1 × 10^10^ CFU/mL, 1 mL). Fibrosis markers were significantly reduced in the IUA + LAC group relative to the IUA group, supporting an antifibrotic effect in vivo. Similarly, ERK1/2 inhibition decreased fibrosis markers, indicating that ERK1/2 blockade alleviates fibrotic responses. Conversely, ERK1/2 activation increased fibrosis markers despite Lactobacillus administration, suggesting that ERK1/2 activation counteracts probiotic-mediated protection (**Fig. 5C–D**).

### 3.6 Characterization of major Lactobacillus delbrueckii–associated metabolites

To integrate microbiome and metabolomics findings, correlation analyses were performed between differential taxa and differential metabolites. *Lactobacillus* abundance showed a positive correlation with 3-HAA levels (**Fig. 6A**), supporting an association between probiotic-related microbial enrichment and increased 3-HAA.

**Figure 6.**
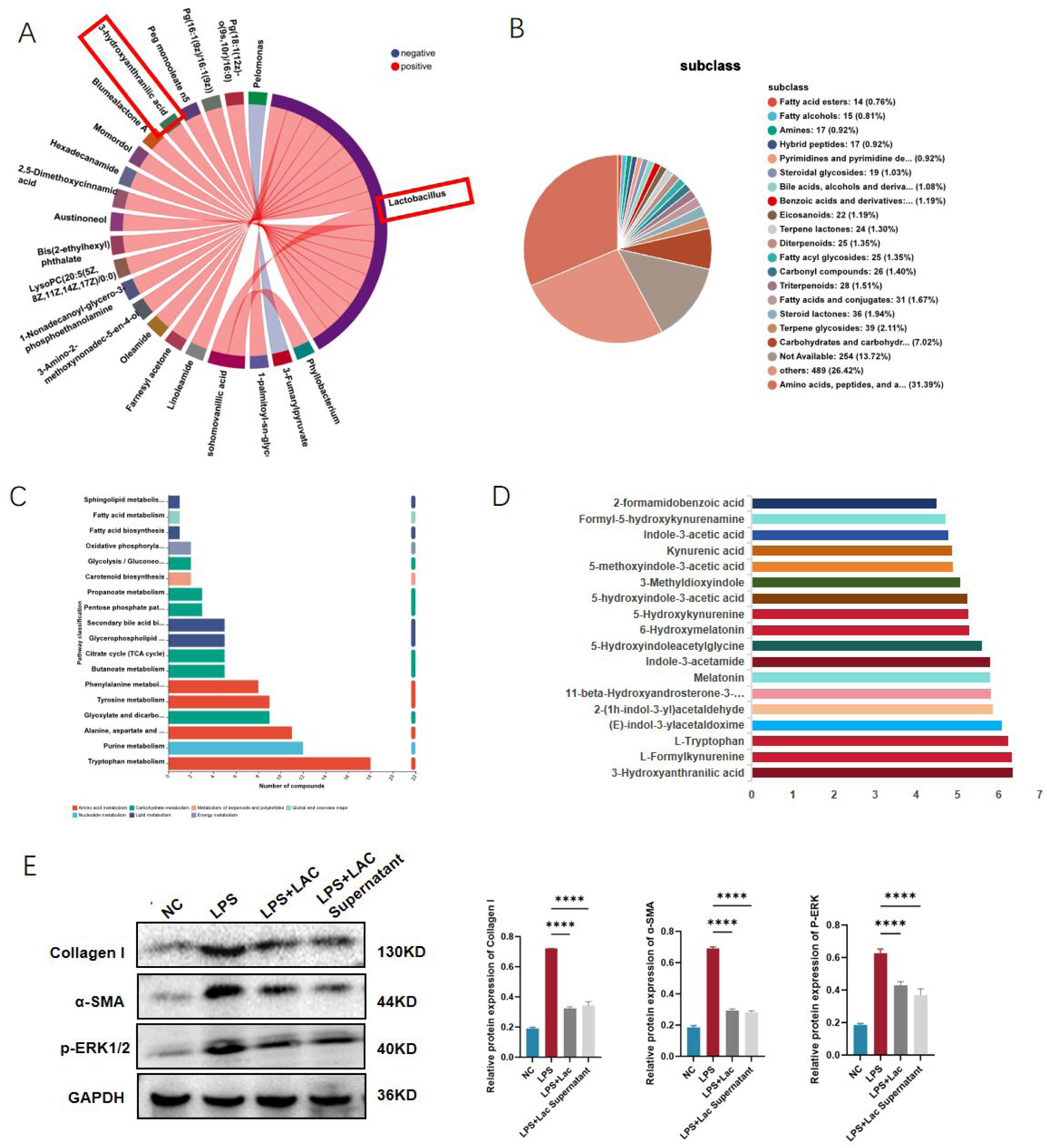
Identification of key metabolites associated with *L. delbrueckii*. (A) Correlation chord diagram between differential taxa and metabolites. (B) Metabolite class distribution. (C) KEGG enrichment for culture supernatant metabolites. (D) Relative abundance of tryptophan pathway metabolites. (E) Effects of L. delbrueckii and its supernatant on collagen I, α-SMA, and p-ERK1/2.

To determine whether *L. delbrueckii* can produce metabolites consistent with this signature, culture supernatants were analyzed by LC–MS. A total of 2554 metabolites were detected. Amino acid–related metabolites accounted for the largest fraction (31.39%), exceeding carbon compounds (7.02%) and terpenoids (2.11%) (**Fig. 6B**). KEGG enrichment revealed significant representation of tryptophan metabolism (**Fig. 6C**). Among detected metabolites within the tryptophan metabolism pathway, 3-HAA showed relatively high abundance compared with other pathway intermediates (**Fig. 6D**).

We next compared antifibrotic effects of direct bacterial addition versus culture supernatant exposure. There was no substantial difference between these approaches, and treatment with 20% *L. delbrueckii* supernatant significantly suppressed fibrotic marker expression (**Fig. 6E**). These findings suggest that secreted metabolites are major contributors to the antifibrotic activity of *L. delbrueckii*, consistent with a microbe–metabolite mechanism.

### 3.7 Validation of 3-HAA as an antifibrotic mediator in cells and mice

To test whether 3-HAA is sufficient to reproduce the signaling and phenotypic effects of probiotic treatment, IHESCs were treated with 3-HAA after LPS induction. At 100 μM, 3-HAA reduced ERK1/2 phosphorylation to a degree comparable with *L. delbrueckii*. Specifically, p-ERK1/2 was significantly decreased in the LPS + 3-HAA group, and fibrotic markers (TGF-β1, α-SMA, collagen I) were also downregulated (**Fig. 7A–B**). These results support a direct antifibrotic role of 3-HAA in the stromal cell model and place ERK1/2 phosphorylation downstream of 3-HAA exposure.

**Figure 7.**
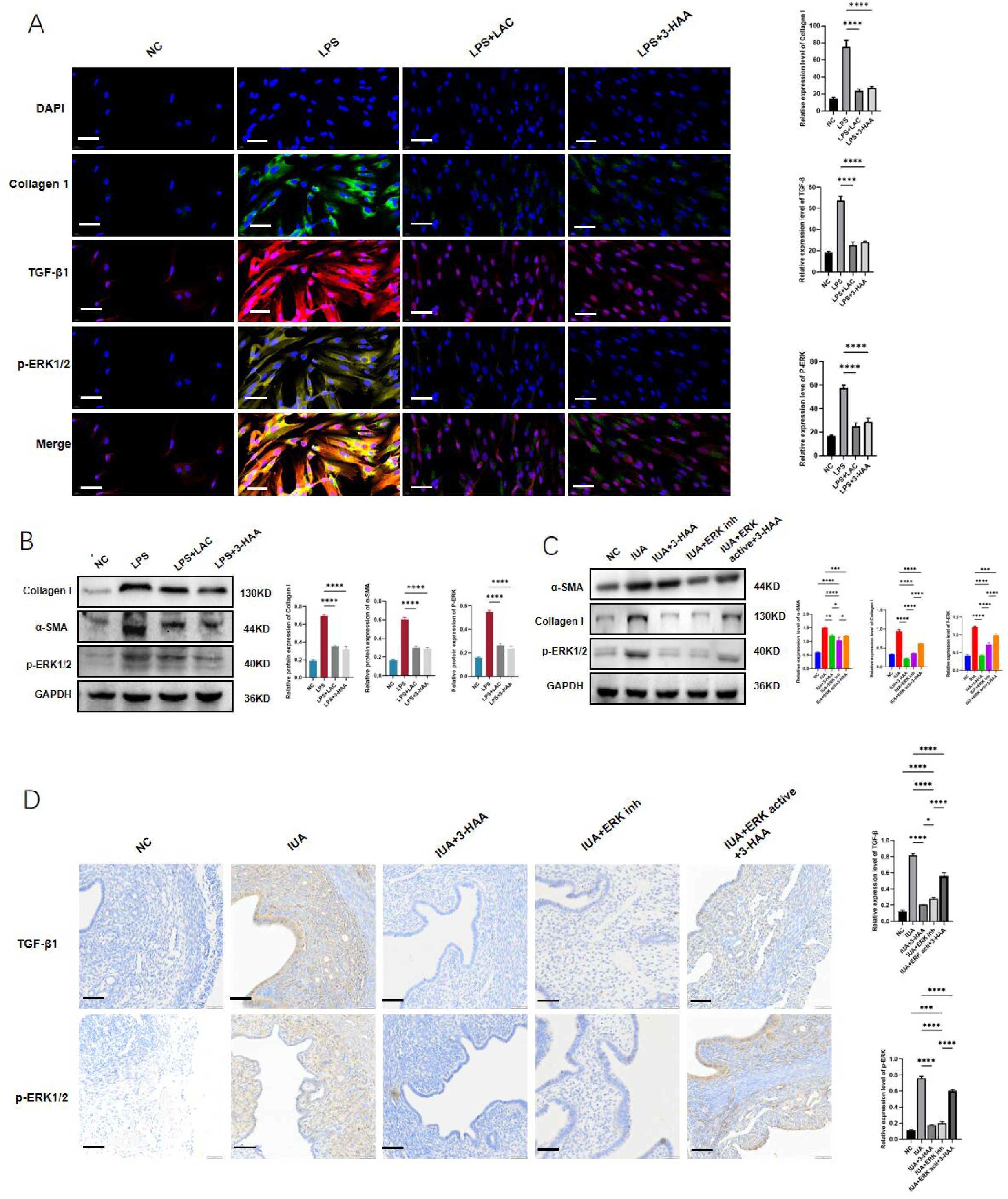
Functional validation of 3-HAA. (A) Immunofluorescence of collagen I, TGF-β1, and p-ERK1/2 after 3-HAA treatment (scale bar = 20 μm). (B) Western blot of collagen I, α-SMA, and p-ERK1/2 after 3-HAA treatment. (C) Western blot of α-SMA, collagen I, and p-ERK1/2 in mouse groups. (D) Immunohistochemistry of TGF-β1 and p-ERK1/2 (scale bar = 50 μm).

In the mouse model, fibrosis markers and p-ERK1/2 expression were lower in the IUA + 3-HAA group than in IUA controls, indicating that 3-HAA also exhibits in vivo antifibrotic activity. ERK1/2 activation increased fibrotic markers and p-ERK1/2 despite 3-HAA treatment, indicating that ERK1/2 activation counteracts 3-HAA–mediated protection (**Fig. 7C–D**). Together, these findings identify 3-HAA as a metabolite mediator capable of suppressing ERK1/2 activation and reducing fibrosis in IUA-relevant models.

## 4. Discussion

Intrauterine adhesions (IUA), commonly referred to as Asherman’s syndrome, represent a significant clinical challenge. The condition, characterized by the formation of fibrous bands within the uterine cavity, leads to various reproductive complications, including infertility, menstrual disturbances, and recurrent pregnancy loss.The increasing prevalence of IUA, particularly in patients with a history of uterine trauma or surgery, underscores the importance of improving diagnostic and therapeutic approaches. Current treatment options, such as hysteroscopic lysis of adhesions combined with hormonal therapy, have yielded suboptimal results, with a high risk of adhesion recurrence and incomplete restoration of endometrial function. Therefore, there is an urgent need for innovative strategies that address the underlying mechanisms of fibrosis and facilitate the repair of the endometrium. This study aimed to explore the therapeutic potential of *Lactobacillus delbrueckii* in modulating the fibrotic processes associated with IUA, focusing on its ability to inhibit the ERK1/2 signaling pathway. By combining clinical data, in vitro models, and multi-omics analysis, we provided new insights into the therapeutic potential of *L. delbrueckii* in regulating uterine fibrosis. The findings presented herein not only contribute to the existing body of knowledge regarding the microbiome’s influence on reproductive health but also pave the way for targeted discussions on the implications of Lactobacillus delbrueckii in clinical practice and future research endeavors.

The present study elucidates the molecular mechanism by which Lactobacillus delbrueckii modulates the ERK1/2 signaling pathway to attenuate uterine fibrosis. Our findings demonstrate that L. delbrueckii significantly reduces the phosphorylation levels of ERK1/2 in LPS-induced human endometrial stromal cells, concomitantly suppressing the expression of fibrogenic markers such as TGF-β1, α-SMA, and Collagen I. This result is in agreement with previous research, where ERK1/2 inhibition in models of cardiac and renal fibrosis has shown that blocking this pathway can reduce myofibroblast activation and matrix deposition^[10–12]^. Specifically, the attenuation of p-ERK levels resulting from *Lactobacillus* intervention mirrors the antifibrotic effects observed with direct ERK1/2 inhibition in models of cardiac and renal fibrosis, where blockade of this pathway leads to diminished myofibroblast activation and reduced extracellular matrix deposition^[13,14]^. Notably, our data extend these insights to the context of uterine fibrosis, demonstrating that probiotic modulation of ERK1/2 signaling is sufficient to downregulate the expression of canonical fibrotic markers such as TGF-β1 and α-SMA. Unlike prior studies that have primarily focused on pharmacologic inhibitors or genetic modulation of MAPK components^[15]^, our results reveal that microbial-derived metabolites or signaling molecules may exert comparable regulatory effects, suggesting a unique host-microbe crosstalk at the level of cellular signaling cascades. While the precise molecular intermediates remain to be fully elucidated, prior investigations have implicated both direct receptor-mediated effects and alterations in local cytokine milieu as potential mechanisms by which probiotics could modulate MAPK activity^[16,17]^. Thus, our findings not only corroborate the centrality of ERK1/2 in fibrotic pathogenesis but also introduce a novel mechanism by which probiotics may interface with host intracellular signaling networks.

Building upon the molecular pathway findings, our analysis of protein expression patterns in mouse endometrial tissue revealed a pronounced reduction in key fibrosis-associated proteins, notably TGF-β1 and α-SMA, following Lactobacillus delbrueckii treatment. This protein-level modulation is consistent with existing evidence from other fibrotic models, wherein inhibition of ERK1/2 signaling leads to downregulation of collagen isoforms and smooth muscle actin, thereby limiting matrix accumulation and myofibroblast transdifferentiation^[18]^. However, whereas pharmacological or genetic interventions in previous work have often resulted in off-target effects or incomplete suppression of fibrotic markers, the probiotic approach employed here appears to achieve a more selective and robust attenuation of fibrosis-related protein expression. It is noteworthy that similar reductions in TGF-β1 and α-SMA have been reported in the context of cardiac and renal fibrosis with ERK1/2 inhibition, supporting the translatability of our findings across tissue types^[19]^. Nevertheless, our data diverge from certain studies where suppression of ERK1/2 activity did not consistently lead to reduced collagen expression, highlighting potential tissue-specific regulatory mechanisms or compensatory pathways that may be differentially engaged depending on the fibrotic niche^[20]^. Collectively, these results underscore the capacity of Lactobacillus delbrueckii to modulate the fibrotic protein landscape via mechanisms that are both convergent with and distinct from established antifibrotic strategies.

In the realm of metabolomic alterations, our study identified a distinct elevation of 3-hydroxyanthranilic acid (3-HAA) in the Lactobacillus delbrueckii-treated cohort, a metabolite previously recognized for its immunomodulatory and anti-inflammatory properties. While the role of 3-HAA in uterine fibrosis has not been widely characterized, analogous shifts in tryptophan-derived metabolites have been implicated in the regulation of immune responses and fibrosis resolution in other organ systems^[21]^. This increase in 3-HAA is particularly intriguing in light of its potential to inhibit key inflammatory mediators and to modulate immune cell differentiation, effects that could synergize with ERK1/2 pathway suppression to curtail fibrotic progression. It is notable that previous probiotic studies have rarely linked microbial administration to specific host metabolite changes with direct relevance to antifibrotic outcomes, marking this as an area of mechanistic novelty. Moreover, these metabolic shifts may represent a broader class of probiotic-host interactions, wherein bacterial metabolites serve as systemic signaling molecules capable of reprogramming host cell fate and matrix dynamics^[22]^. This metabolomic signature thus offers a new perspective on how probiotics can exert tissue-specific therapeutic effects beyond local microbiota modulation.

The metabolomic analysis in our study also revealed that 3-HAA is closely linked to microbial changes within the uterine cavity. Lactobacillus administration led to increased microbial diversity and the enrichment of beneficial microbial species, which was correlated with the reduction of fibrotic markers. These findings support the idea that probiotics, by reshaping the microbial ecosystem, can create a more favorable environment for endometrial repair. This result is consistent with previous studies that have shown that a more diverse microbiome, particularly one enriched with Lactobacillus species, is associated with better reproductive outcomes and reduced gynecological pathologies^[23]^. Notably, our findings demonstrate that targeted probiotic supplementation can remodel the uterine microbial ecosystem in a manner that correlates with attenuation of fibrosis, an association that has been suggested but not mechanistically established in prior studies^[24]^. Furthermore, while intravaginal or intrauterine administration of probiotics has been proposed as a means to restore microbial homeostasis and counteract pathogenic shifts, empirical evidence for direct impact on fibrotic disease processes has been limited; our data provide a concrete link between microbiota composition and host tissue remodeling. In contrast to earlier studies that often relied on observational correlations without interventional validation^[25]^, the present results substantiate a causal relationship, highlighting the therapeutic potential of microbial interventions in modulating both microbial ecology and fibrotic pathophysiology.

Despite the promising results, several limitations should be acknowledged.First, the relatively small sample size limits the generalizability of the findings. A larger cohort would provide more robust data and reduce the potential for variability in individual responses to *L. delbrueckii* treatment. Second, the lack of long-term follow-up data restricts our ability to assess the durability of the therapeutic effects. Future studies should investigate whether the observed improvements in fibrosis markers persist over time and whether *L. delbrueckii* can prevent adhesion recurrence in the long term. Additionally, batch effects across the different datasets analyzed in this study could introduce variability, which may complicate the interpretation of the results. These issues could be mitigated by incorporating standardized protocols and increasing sample sizes in future research.

In conclusion, this study provides new insights into the therapeutic potential of *Lactobacillus delbrueckii* for managing IUA-associated fibrosis. By inhibiting the ERK1/2 pathway and promoting favorable microbial changes, *L. delbrueckii* offers a novel, non-invasive approach for improving uterine health and fertility outcomes. These findings have important clinical implications, suggesting that probiotics could serve as a complementary treatment for IUA, especially in patients who are at high risk of recurrence following surgical interventions. Future research should focus on validating these findings in larger, multicenter trials and exploring the long-term efficacy and safety of *L. delbrueckii* in preventing IUA recurrence and improving reproductive health.

## Declarations

### Ethics approval and consent to participate

This study was approved by the Ethics Committee of Chongqing Health Center for Women and Children (Approval No. 2021044). All participants provided written informed consent prior to sample collection.

### Availability of data and materials

All datasets supporting the conclusions of this study are included in the article and its supplementary files.

## Acknowledgment

We thank all participants and colleagues who contributed to this study and to manuscript preparation.

